# Glaciation history and geography shape genetic diversity of Lewis flax (*Linum lewisii* Pursh.) across North America

**DOI:** 10.1101/2025.08.18.670546

**Authors:** Peter Innes, Zach Marcus, Brent S. Hulke, Nolan C. Kane

**Author notes:** Corresponding author Peter Innes 1900 Pleasant Street 334 UCB Boulder, CO 80309.

## Abstract

Factors shaping genetic variation at continental scales include geography, glacial cycles, and genetic constraints. We examine how such factors influenced diversification in *Linum lewisii* Pursh. (Lewis flax), a perennial forb which colonized and spread across North America in the last few million years, accompanied by self-incompatibility and distyly loss. We used whole-genome analyses to quantify population structure and diversity across the landscape, estimate population histories, and test expectations about the shift to self-compatibility; we examine how understanding the evolutionary history of this broadly-adapted species informs ecological restoration and neodomestication efforts. We show that Lewis flax consists of four genetic clusters that diverged during the Last Glacial Period or earlier, with mountainous regions acting as historical barriers, contemporary dispersal corridors and contact zones. The four lineages within Lewis Flax are differentiated but do not correspond to existing taxonomic treatments. Our results support the hypothesis of *refugia within refugia*, the presence of population substructure within southern refugia. Genomic data further indicate Lewis flax is predominantly outcrossing despite its self-compatibility, although postglacial range expansions may have involved shifting towards increased selfing. The mixed mating system of Lewis flax thus interacts with glacial history and geographical features to shape diversification across the continent.

## Introduction

Glacial cycles have shaped global patterns of population genetic diversity through profound alteration of climate, continental landscapes, and demographic trajectories (Fonseca et al., 2023; Hewitt, 2004). During cold periods, many Northern Hemisphere temperate plant and animal species experienced range contractions into climatically suitable refugia, followed by postglacial expansion in warmer eras (Comes & Kadereit, 1998; Hewitt, 2000; Hewitt, 1996). Because genetic variation tends to accumulate in populations that persist through time, while range expansions typically involve genetic bottlenecks, there is an expectation of higher diversity in refugial populations relative to those in newly colonized territory (Comes & Kadereit, 1998). Furthermore, if a species inhabited multiple distinct glacial refugia, genetic divergence between the separated populations is likely to occur due to genetic drift and local adaptation, and genetic structuring may occur along the expansion path as well (Hewitt, 1996). The early and middle Pleistocene saw frequent speciation in this fashion (Kadereit & Abbott, 2021; Lovette, 2005). For most species, however, little is known about the specific impacts of Pleistocene climatic oscillations on intraspecific diversification (Ostrow et al., 2025). An intuitive hypothesis is that late Pleistocene glaciation (e.g. the Last Glacial Period) was an underlying cause of population-level divergence, which is assumed to occur at shallower time scales relative to speciation. This effect likely persists to this day in patterns of genetic variation across extant populations of many species.

In the classic paradigm for western North America, major glacial refugia existed in two regions: south of the continental ice sheet and north, in Beringia (Shafer et al., 2010). More recent studies emphasize historical subdivision and population isolation within these broad regions (i.e. “refugia within refugia”, as well as the existence of alternative “cryptic" northern refugia (Beatty & Provan, 2010; Hewitt, 2004; Provan & Bennett, 2008; Shafer et al., 2010). Mountain ranges including the Rocky Mountains, Coast Range, and Cascade–Sierra chain also factor heavily in the phylogeography of this region: examples include discontinuities between different ranges (Brunsfeld et al., 2001), north–south differentiation along mountain chains (Jaramillo-Correa et al., 2009), mountains acting as zones of secondary contact between divergent taxa (Remington, 1968; Swenson & Howard, 2005), and mountain chains acting as dispersal corridors (Shafer et al., 2010). These patterns deserve continued examination in light of the fact that, overall, many species in North America exhibit unique histories regarding the refugia and geographic features that have shaped their contemporary diversity (Soltis et al., 2006).

For plant species, reproductive strategy, i.e. mating system, is an additional factor that may interact with climatic history and geography in shaping population genetic diversification across the landscape (Koski et al., 2019; Urquhart-Cronish et al., 2026). For instance, self-compatibility can facilitate range expansion by improving reproductive assurance at distribution edges where mate availability is limited (Baker, 1955); this could allow for rapid expansion following major environmental changes like glacial abatement. Selfing is also expected to lead to stronger population structure due to reduced pollen dispersal and gene flow with neighboring plants and populations (Wright et al., 2013); while selfing is often hypothesized as an evolutionary dead-end, species with mixed mating systems (i.e. a combination of outcrossing and selfing) may exhibit increased diversification rates owing to the benefits of both strategies (Goodwillie et al., 2005; Wright et al., 2013). How species with mixed mating systems respond to dramatic environmental shifts is thus of particular interest.

Here, we explore how late Pleistocene glaciation and major landscape features in western North America have combined to influence population genetic diversification in *Linum lewisii* Pursh., commonly known as Lewis flax or prairie flax, a self-compatible perennial forb with a distribution spanning western North America from Alaska to northern Mexico, with some isolated populations around Hudson Bay in Canada (Fig. S1; Mosquin, 1971). Lewis flax is frequently included in native ecosystem restoration seed mixes, as it is one of only a handful of native forbs for which seed is commercially produced (Jones, 2019). There is also an ongoing effort to domesticate Lewis flax as a new perennial oilseed crop given its similarities to common flax (*Linum usitatissimum*), the source of flaxseed, linseed oil, and linen (Pull et al., 2023). There are two other blue flax species native to North America: *Linum pratense* (Norton) Small, which has a smaller range, overlapping with the southeastern extent of *L. lewisii*, and the lesser known *Linum rzedowskii* Arreguín, which has a very restricted endemic range in central Mexico and is critically endangered (González-Velasco et al., 2022). All other close relatives of Lewis flax inhabit Eurasia. The ancestor of Lewis flax is thought to have colonized North America via Beringia during its split from Eurasian populations 3.3–3.8 Mya (Harris, 1968; McDill et al., 2009).

During or preceding this colonization, a notable evolutionary shift occurred: Lewis flax transitioned from self-incompatibility to self-compatibility (McDill et al., 2009; Ruiz-Martín et al., 2018). In the *Linum* genus, self-incompatibility is tied to heterostyly (i.e. distyly), an iconic floral polymorphism defined by the presence of (typically) two genetically determined mating types with distinct floral morphology (Darwin, 1863). Heterostyly is believed to be the ancestral state of the genus but has been lost several times independently, leading to homostyly, in which plants exhibit a single, self-compatible floral morph (Maguilla et al., 2021; McDill et al., 2009). Recent findings in yellow flax and in the genus *Primula* have shown how distyly loss results in strong selfing (Gutiérrez-Valencia et al., 2024; Mora-Carrera et al., 2023). Lewis flax is homostylous and self-compatible, although its propensity for selfing versus outcrossing is unclear. A previous pollination experiment suggests it does not autonomously self (Kearns & Inouye, 1994), plausibly due to its approach herkogamy, defined as the spatial separation of anthers and stigmas, specifically with the latter extending beyond the former (Webb & Lloyd, 1986). (Here we note that we use “homostyly” in the general sense, to refer to species with monomorphic floral morphology following loss of heterostyly regardless of relative positions of anthers and stigmas, whereas elsewhere it is sometimes used to describe monomorphism specifically with anthers and stigmas at the same height). Approach herkogamy is uncommon among homostylous flaxes, which typically have anthers and stigmas at the same height (Ruiz-Martín et al., 2018). A separate survey of Lewis flax herbarium material reported a reduction in the degree of herkogamy in some parts of its range (Mosquin, 1971), suggesting possible variation in selfing rates as well. Although existing taxonomic treatments have yet to be confirmed by genetic analysis, herkogamy is thought to vary among some of the named lineages (Flora of North America Editorial Committee, 2016).

We sought to answer the following major question: How have historical glaciation and major landscape features influenced population diversification in Lewis flax? Given the broad distribution of Lewis flax, we tested the “complexity” hypothesis *sensu* Shafer *et al*. (Shafer et al., 2010), which extends the Beringia–southern paradigm to incorporate historic substructure i.e. *refugia within refugia* and the possibility of cryptic northern refugia. Additionally, we examined the extent to which loss of distyly in Lewis flax has resulted in selfing, and whether inbreeding differs across its diverse geographical and ecological range. We hypothesized that, despite its shift to self-compatibility, herkogamy has precluded the evolution of strong selfing in Lewis flax. We used whole genome sequencing to characterize the major genetic lineages of Lewis flax across the western North American landscape and how they relate to current infraspecific taxonomy. We inferred population size trajectories and divergence times over the past million years, and we looked for genomic signatures of selfing in terms of homozygosity, linkage disequilibrium, and effective population size. Lastly, we discuss how our findings might be applied to the use of Lewis flax in restoration and agricultural settings. Our study uncovers biogeographic patterns and their causes at the continental scale and illuminates population genetic trends following an unusual case of distyly loss with herkogamy.

## Materials and methods

### Plant material

Our initial sampling combined 172 Lewis flax genotypes from seed banks (USDA National Plant Germplasm System and Plant Gene Resources of Canada, *n* = 34), herbaria (*n* = 66), personal seed collections (*n* = 70), and the National Center for Biotechnology Information Sequence Read Archive (NCBI SRA, *n* = 2) (Table S1). Sampling was conducted opportunistically due to limited resources and travel prohibitions during the COVID-19 pandemic. Most geographic locations are represented by a single genotype (Figure S2), with the main exception being that we had dense sampling in Boulder County, Colorado (*n* = 32 across 13 localities).

Seed bank accessions and personal collections were grown from seed either in a greenhouse setting at University of Colorado, Boulder or in field settings at North Dakota State University and University of Minnesota, Twin Cities. We sampled herbarium specimens from the University of Colorado Herbarium (COLO), the Rocky Mountain Herbarium at University of Wyoming (RM), the University of Manitoba Vascular Plant Herbarium (WIN), the Arizona–Sonora Desert Museum (ASDM), and the Desert Botanical Garden (DES). Acronyms follow the Index Herbariorum database. We selected specimens that filled geographic gaps among our seed collections, were recently collected, appeared in good condition, and had enough leaf material to minimize the impact of destructive sampling (we took leaf tissue from fragment packs associated with the specimen whenever possible). Our study aims did not have a temporal component beyond historical inference made from contemporary samples, and thus we prioritized specimens we thought could yield high quality DNA without using intensive ancient DNA protocols.

Of the two *L. lewisii* whole genome shotgun (WGS) samples retrieved from the SRA, Llw01 was from a genotype of unknown provenance (You et al., 2018); the other, TROM V 135817, was from an Arctic University Museum of Norway herbarium specimen collected in Alaska (Alsos et al., 2020). Eight samples lacked geographic coordinates or detailed locality information (Table S1) and thus were excluded from spatial analyses and map visualizations.

### DNA extraction, library preparation, and sequencing

Extractions and library preparations were performed separately for non-herbarium samples versus herbarium samples. For the former, we extracted DNA from fresh, lyophilized, or silica-dried leaf tissue using QIAGEN DNeasy Plant Mini kits. Libraries for Illumina whole genome sequencing were prepared with the 96-Plex kit from TWIST Biosciences following the manufacturer’s protocol and using an even ratio of the Low and High GC Primer A. DNA amounts across samples were equalized for library input by dilution based on Qubit fluorometer readings. Samples were spread across two TWIST 96-Plex libraries. Library quality control and sequencing was performed by Novogene. The pooled Twist libraries were each sequenced on a single lane of a Nova-Seq 6000 S4 flow cell using 150 bp paired-end reads.

Herbarium sample DNA extractions were performed several weeks to months after the non-herbarium sample extractions but in the same lab space at University of Colorado, Boulder. Equipment and surfaces were thoroughly cleaned with 10% bleach in order to safeguard against contamination; we also treated plastic consumables with UV light prior to extractions. We used a modified QIAGEN DNeasy Plant Mini Kit extraction protocol optimized for herbarium specimen leaf tissue (Marinček et al., 2022). We assessed herbarium DNA quantity and quality with flourometer (Qubit), spectrophotometer (Nanodrop), and gel electrophoresis methods. Herbarium DNA samples passing this quality control step had concentrations ranging from 2–34 ng/µL (mean 13 ng/µL), a 260/280 ratio ranging from 1.6–2.8, and fragments mostly 500 bp or larger based on gel results. A preliminary attempt to include herbarium DNA samples in a pooled Twist 96-Plex library alongside non-herbarium samples resulted in very low amounts of sequence data for the herbarium samples, so we took a different approach. Illumina whole genome libraries were prepared for herbarium DNA samples of sufficient quality using the Illumina DNA Prep workflow (formerly Nextera DNA Flex), which includes an initial fragmentation step. The libraries were sequenced on a single lane of an Illumina NovaSeq X Plus 10B flow cell using 150 bp paired-end reads. Herbarium WGS library preparation and sequencing were performed by SNPsaurus.

Among the 104 non-herbarium samples that were sequenced, we excluded four of five samples that were grown from seed from the same mother plant in Boulder County, Colorado, due to their sibling relationship (Table S1). We also excluded eight of 66 herbarium samples from analysis of the nuclear genome due to low depth (four had depth less than 0.25x and another four had depth less than 0.75x).

### DNA sequence processing, alignment, variant calling, and filtering

Raw sequence data were quality trimmed using fastp with default settings (Chen et al., 2018) and aligned to the chromosome-scale *L. lewisii* Maple Grove reference genome [GenBank: GCA_034768395.1 (Innes et al., 2023)] with bwa-mem2 (Vasimuddin et al., 2019). We marked duplicate reads with Picard MarkDuplicates and used mosdepth (Pedersen & Quinlan, 2018) to calculate average sequencing depth for each sample.

We called variants with bcftools mpileup and bcftools call (Danecek et al., 2021). We used GNU parallel (Tange, 2023) to process the nine chromosomes of the Lewis flax genome in parallel and generate VCF files for each chromosome that contained both variant and invariant sites. We excluded indels while filtering SNPs and invariant sites with the following site-level thresholds using bcftools: we kept sites with minimum average mapping quality (MQ) 40, maximum missingness (F_MISSING) 25%, and mean depth [MEAN(FMT/DP)] between 2 and 15. For SNPs we additionally kept only bi-allelic sites with quality (QUAL) of at least 40. The maximum mean depth threshold (15) was slightly less than three times the median of per-sample average depth values and was meant as an initial filter for spurious calls in repetitive or highly paralogous regions. We used two additional filters to limit potential biases resulting from paralogy and repetitiveness in the Lewis flax genome, approximately 70% of which is repetitive (Innes et al., 2023). First, using mosdepth, we calculated depth averaged within 500 bp windows across the genome; for each sample, windows with depth greater than 1.5x the within-sample median were defined as high depth regions; we then defined a set of shared high depth regions as those high depth regions identified in at least half of all samples. Second, we used GenMap (Pockrandt et al., 2020) to calculate mappability across the Lewis flax reference genome with a k-mer length of 150 bp and up to three errors (parameters: -K 150 -E 3); low mappability regions were defined as those matching three or more places in the genome. We merged these regions with the shared high depth regions using bedtools. The resulting sites, comprising 262,564,216 bp, around 43% of the length of the nine reference genome chromosomes, were further excluded from our VCF. Filtered variant and invariant sites were then combined back into a filtered “all-sites” VCF containing 123,583,335 invariant sites and 16,218,244 biallelic SNPs, of which 5,417,581 were singletons (i.e. had a minor allele count of 1).

### Integrity of herbarium specimen DNA

We were cautious about including herbarium specimens alongside contemporary samples in our analyses, due to potential batch effects that could arise from DNA degradation or the use of different library preparation kits. We used mapDamage2 (Jónsson et al., 2013) to quantify damage patterns in our herbarium DNA and examined the frequency of cytosine deamination (C to T transitions) at the ends of reads, a hallmark of post-mortem degradation. We also looked for batch effects in our subsequent analyses of population structure and diversity simply by comparing technical versus geographic groupings.

### Population structure

For analyses of population structure, we used a minimum minor allele frequency threshold of 0.025 and also pruned SNPs based on linkage disequilibrium (LD) using PLINK v1.90b7.1 (Chang et al., 2015) with a window size of 10 kb, a step size of 1 SNP, and a maximum *r*^2^ threshold of 0.5 (`plink --indep-pairwise 10kb 1 0.5`). This resulted in 1,349,461 SNPs. We performed model-based ancestry inference with ADMIXTURE (Alexander et al., 2009) for values of *k* ranging from 1 to 10 and using 5-fold cross validation to determine the most likely value of *k*. Principal component analysis (PCA) was performed with EMU (Meisner et al., 2021), to better account for non-random missingness stemming from variation in sequencing depth among samples; we modeled and extracted 10 eigenvectors. Results from both ADMIXTURE and PCA suggested the *L. lewisii* samples included here could be assigned to four main clusters, which we hereafter refer to as the Northern (N), Central (C), Southwestern (SW) and Southeastern (SE) populations, based on the ancestry coefficient with relative majority. We subsequently used ancestral population allele frequencies output by ADMIXTURE at *k* = 4 to construct a neighbor-joining tree based on Euclidean distance, using the R package ape v5.8-1 (Paradis & Schliep, 2019).

We investigated the impact of spatial bias in our sampling (particularly the dense sampling in Boulder County, Colorado) by excluding 20 of the 28 Boulder County samples in the full cohort and again running ADMIXTURE and PCA as described above. Prior to analysis we re-filtered the sub-sampled VCF with the same missingness, minor allele frequency, and LD thresholds as described previously. We refer to this dataset as the “reduced cohort”.

We used FEEMS (Marcus et al., 2021) to visualize patterns of Lewis flax effective migration and gain additional insight into its spatial genetic structure. For this analysis we excluded nine samples without geographic coordinates as well as the sample from West Virginia (Ames 34946) and subsequently applied the same minor allele frequency filter and LD pruning as before. Missing genotypes were imputed with Beagle v5.4 (Browning et al., 2018) prior to LD pruning. Migration was estimated across a spatial grid of triangular cells, each with an area of approximately 6,200 square kilometers, which we cropped to the rough outline of our sample area. We used leave-one-out cross-validation to select the best value of the tuning parameter, *λ*, following the FEEMS documentation. In a second FEEMS run, we excluded the 13 samples in the formerly glaciated portion of North America, in order to gain better resolution of migration patterns for the core portion of the Lewis flax range.

To further examine geographic trends in population structure, we examined correlations between genetic principal components and latitude and longitude using the non-parametric coefficient Kendall’s *τ*. We also performed a Mantel test of isolation-by-distance based on Hamming genetic distances between each pair of samples, calculated with PLINK, as well as pairwise geographic distances (km), calculated with the Vincenty Ellipsoid method in the geosphere R package (Hijmans, 2026).

### Genetic diversity, differentiation, and inbreeding

We estimated pairwise nucleotide diversity (*π*) of each population and absolute genetic divergence (*D_xy_*) among populations in 10 kb non-overlapping windows using pixy v1.2.10.beta2 (Korunes & Samuk, 2021) and the filtered all-sites VCF without singleton SNPs. Genome-wide average *π* and *D_xy_* for each population was calculated as the sum of counts of differences (across all 10 kb windows) divided by the sum of counts of comparisons, as recommended in the documentation for pixy. We generated continuous maps of *π* and observed heterozygosity with the wingen R package and used its rarefaction approach to account for differences in sampling effort across the landscape (Bishop et al., 2023). Genome-wide average *F_ST_* for each population pair was calculated across the same SNPs as for estimation of *π* and *D_xy_*, using the function average_hudson_fst() from the python library scikit-allel v1.3.7 (Miles et al., 2021).

We used PLINK to estimate the inbreeding coefficient (*F*) for each sample, based on the same SNPs as used for the diversity and divergence metrics, however, we subset our VCF by genetic cluster in order to limit upward bias due to the Wahlund effect (Hemstrom et al., 2024). To look for evidence of increased selfing at northern latitudes, we fit a basic linear model for *F* ∼ *latitude*.

LD decay curves were generated with PopLDdecay (Zhang et al., 2019), using the filtered SNPs and a minimum minor allele frequency threshold of 0.025.

### Demographic inference

We inferred population size histories and divergence times of the four genetic clusters with SMC++ (Terhorst et al., 2017). SMC++ assumes a “clean split" model to estimate divergence times, in which gene flow ceases at the time of the split. In cases where post-split gene flow does occur, split times are likely to be underestimated, i.e. biased towards more recent times. We took precautions against such bias by limiting our demographic inference to only those plants with minimal admixture (*q* ≥ 0.9), following Coe et al. (2023) and Gaccione et al. (2025). We selected SMC++ despite this potential limitation because it otherwise is an excellent fit for the size and characteristics of our dataset, comprising dozens of unphased whole genomes sampled approximately randomly across the species’ range; SMC++ is robust to error caused by continuous population structure across the landscape (Battey et al., 2020). We further limited this analysis to samples with at least 5x depth. We used SNPs as filtered above for genetic diversity analyses but without a minor allele frequency threshold so as not to alter the site frequency spectrum. We masked the same high depth and low mappability sites as excluded during initial variant filtration. We converted variant data from VCF to SMC++ format using the command smc++ vcf2smc and iterated over five “distinguished individuals” from each population to form a composite likelihood, which we subsequently used for estimation of population size histories (command: smc++ estimate) and split times (command: smc++ split). We used the *Arabidopsis* per-generation mutation rate of 7e-9 (Ossowski et al., 2010) and a generation time of two years given the short-lived perennial habit of Lewis flax. We note that divergence time estimates are sensitive to both mutation rate and generation time, with shorter generations giving estimates of more recent splits. Lewis flax is capable of flowering in the first year, but does not always do so, based on personal observation. Thus, while generation time is an additional source of uncertainty in our demographic modeling, we feel that two years is a better estimate for average age at reproduction in Lewis flax compared to one year. We set beginning and ending time points to 1e3 and 1e6 years ago, respectively. We accounted for uncertainty in population size histories and split times by resampling the input genomic data of each distinguished individual in 5 Mb blocks for 10 bootstrap replicates, following (Coe et al., 2023). For additional insight into demographic history, we calculated genome-wide Tajima’s *D* for each population using pixy v2.0 (Bailey et al., 2025) with the all-sites VCF and no minor allele count filter; we used whole-chromosome windows and values were aggregated across the nine chromosomes to obtain genome-wide estimates.

### Chloroplast phylogenetics

We assembled a complete chloroplast genome for Maple Grove from short-read Omni-C data originally used for reference genome scaffolding (Innes et al., 2023) but which we found to have sufficient plastid genome coverage for assembly. A draft assembly was performed with SPAdes v3.11.1 (Prjibelski et al., 2020) and four resulting plastid scaffolds were manually joined and circularized. The chloroplast genome was included with the nuclear reference genome for alignments described previously. We included *Linum perenne* and *Linum leonii* as outgroups using publicly available whole genome shotgun data [GenBank: PRJNA262458 (You et al., 2018)]. Variants in the chloroplast genome were called using bcftools with ploidy set to 1. We retained only bi-allelic SNPs with QUAL scores of 100 or greater; we also excluded sites within the second copy of the inverted repeat. SNPs in VCF format were converted to phylip format with vcf2phylip.py (Ortiz, 2019). We used iqtree2 for maximum-likelihood phylogenetic inference with ModelFinder and a partition model delineating SNPs in the long single copy, inverted repeat, and short single copy regions (Chernomor et al., 2016; Kalyaanamoorthy et al., 2017; Minh et al., 2020). The model designation was given as -m MFP+ASC+MERGE to account for SNP data ascertainment bias and to allow for potential merging of partitions. Node support was determined with 1000 bootstrap replicates (Hoang et al., 2018). The resulting tree was visualized with ggtree (Yu et al., 2017), and nodes with less than 75 bootstrap support were collapsed.

## Results

### Successful whole-genome sequencing of herbarium DNA

The 160 samples included in the nuclear genome analysis had an overall mean and median sequencing depth of 6.8x and 5.6x, respectively. Herbarium samples (Illumina libraries) ranged from 0.75x to 24x depth, with mean 8.5x and median 5.9x depth. The non-herbarium samples (Twist libraries) ranged from 1.3x to 12x, with mean 5.8x and median 5.6x depth. Percentage of reads mapping to the Maple Grove reference genome was 95%–99% in the non-herbarium samples and 97%–99% among the herbarium samples, suggesting the relative level of endogenous DNA in our herbarium samples was high and contamination was not an issue.

Signs of degradation among the herbarium DNA samples were minimal. The frequency of C to T transitions at the 5’ end of reads was not elevated; nor did it significantly correlate with the age of the specimen (Fig. S3). Sequencing depth was also uncorrelated with specimen age (Fig. S3). We did observe a weak association between herbarium specimen age and DNA library insert size (Fig. S3), but overall, these results support our decision to not treat the herbarium samples separately in our bioinformatic pipelines. The integrity of the herbarium DNA is believable considering we purposefully chose specimens appearing in good condition and which were mostly collected in the past four decades.

### Lewis flax population structure corresponds to four geographic regions

We found the strongest support for *k* = 4 ancestral populations, which generally correspond to the northern, central, southwestern, and southeastern parts of its range (Figs. 1, S4). The Northern population comprised Alaskan and Canadian samples (including the two *L. lewisii* var. *lepagei* samples from Hudson Bay) and reached south—mostly east of the Continental Divide of the Americas—into Montana and the Dakotas, as well as Wyoming (Fig. 1). The Central population had the largest number of samples and tracked the Rocky Mountains—primarily west of the Continental Divide—as well as the Cascade Range in Oregon and into California (Fig. 1).

**Figure 1:**
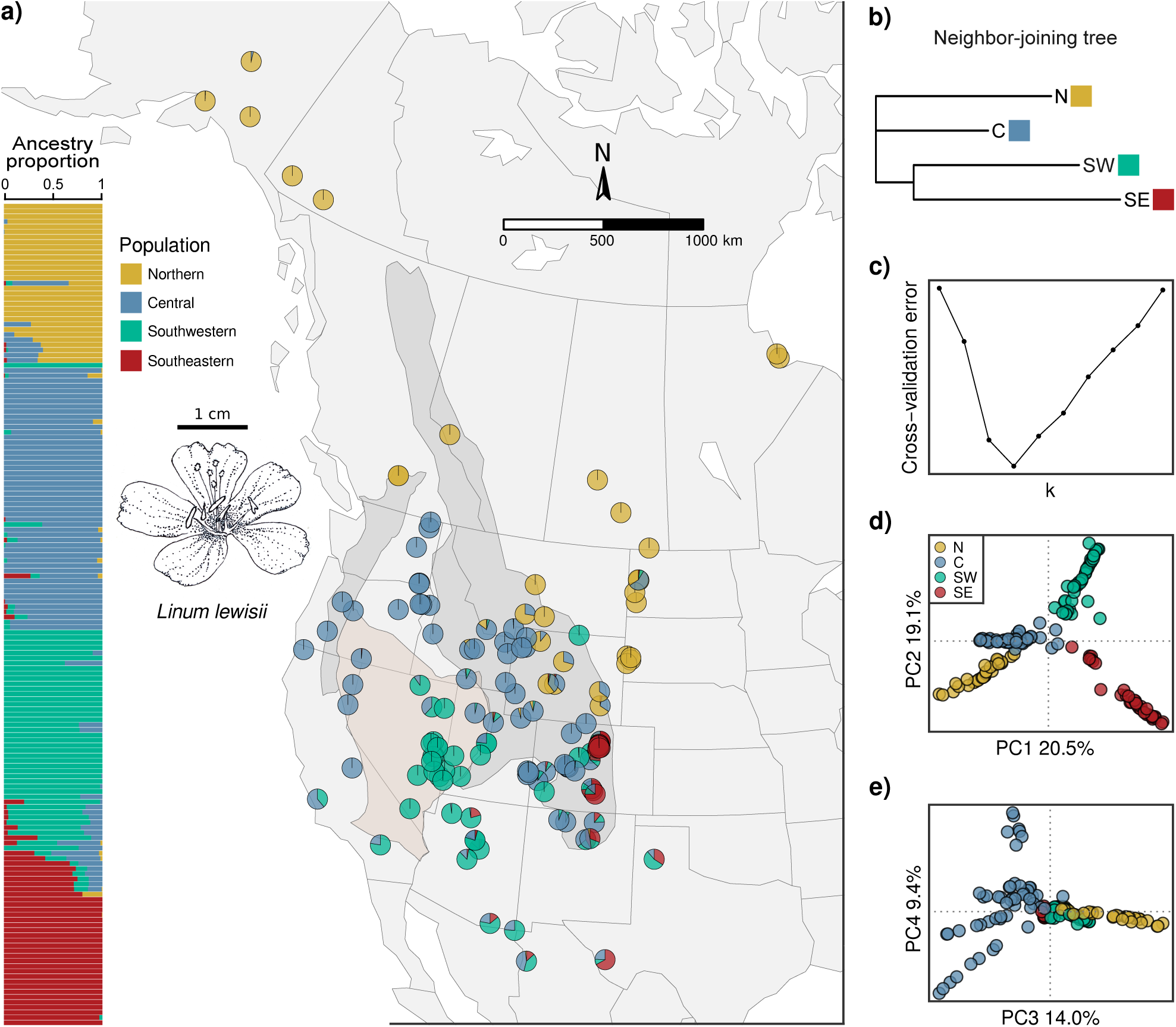
Population genetic structure of Lewis flax across North America inferred from 1,349,461 nuclear SNPs. **a)** ADMIXTURE results and geographic context for *k* = 4. Pie charts represent ancestry coefficients of individual samples. The darker gray shading represents approximate boundaries of the Rocky Mountains and Cascade Range, and the beige shading is the Great Basin, obtained from Natural Earth public map data. For visualization purposes, the Southwestern-type accession Ames 34946 from West Virginia is not shown. The illustration shows a typical flower of *Linum lewisii*, by Karen Jepson-Innes. **b)** Neighbor-joining tree based on inferred ancestral population allele frequencies output by ADMIXUTRE. **c)** Cross-validation errors of ADMIXTURE runs for *k* = 1 through 10. **d–e)** Principal component analysis of SNP variation. Points are individual samples and are colored based on the relative majority ancestry coefficient at*k* = 4

The Southwestern population mainly consisted of samples from the Great Basin in Nevada and Utah, as well as samples from Arizona. Furthermore, the sample from the San Bernardino Mountains in southern California (COLO 539427) and the sample from Sonora, Mexico (ASDM 3338) both had predominantly Southwestern ancestry, while the sample from Chihuahua, Mexico (RM 55736) was more admixed, with a relative majority of Central ancestry. The Maple Grove genotype (Ames 27614), which was used for the reference genome assembly (Innes et al., 2023), had an estimated 99.8% Southwestern ancestry. A few samples with 100% Southwestern ancestry were located far outside the core Southwestern region. These included the sample from a naturalized population in West Virginia (Ames 34946; Table S1, Fig. S2), as well as Ames 34266 (northern Wyoming), and three samples from central Colorado (Guanella Pass High, Homestake Reservoir, and Dow Mountain).

The Southeastern population appeared mainly in the Southern Rockies on the eastern side of the Continental Divide. This population mostly included samples from around Boulder County, Colorado, where our sampling was particularly dense, though admixed samples with varying amounts of Southeastern ancestry occur in New Mexico and Texas and, to a lesser extent, Arizona and northern Mexico. When we thinned our sampling in Boulder County from 20 down to 8 samples, several plants were estimated to have a larger amount of Southeastern ancestry (Fig. S4). For instance, the sample from southwestern Texas (COLO 90125) showed 67% SE ancestry in the analysis of the full cohort but 100% SE ancestry in the context of the reduced cohort (Figs. 1, S4). Further regional population substructure is apparent in the Central population at *k* = 5 and *k* = 6 (Figs. S5, S6). At *k* = 7, the Southwestern population is divided latitudinally (Figs. S5, S6). Smaller geographic clusters in Colorado, Wyoming, and New Mexico appeared in *k* = 8 through *k* = 10 but are difficult to interpret given the low support according to cross-validation error scores (Figs. S5, S6). It is possible that additional sampling, especially towards range margins where our sampling tended to be more sparse, may reveal additional population structure not discovered here. Principal component analysis of SNP variation also indicated four clusters. The first axis (20.5% variation explained) separated the Northern and Central populations from the Southeastern and Southwestern populations and showed a significant correlation with latitude (Kendall’s *τ* = −0.41; *p* < 0.001) (Fig. S7). The second axis (19.1% variation explained) captured additional structure among all four populations (Fig. 1) and was correlated with longitude (Fig. S7). We note that the values of variance explained by each principal component should be interpreted only relative to each other rather than in absolute terms because EMU uses a truncated implementation of singular-value decomposition and therefore does not provide the full eigenspectrum for proper normalization (Meisner et al., 2021). Visualization of PC3 and PC4 revealed more fine-scale population structure within the Central population (Fig. 1), which is consistent with ADMIXTURE results for *k* = 5 and *k* = 6 (Figs. S5, S6).

Geographic and genetic distances among samples showed a moderate and significant correlation via Mantel test (*r* = 0.12, *p* = 0.007; Fig. S7c), consistent with the association between genetic principal component and geographic coordinates.

We examined sequencing batch and variation in sequencing depth among samples as potential sources of technical bias in our inferences about population structure. We did not observe obvious batch effects between herbarium and non-herbarium samples in the PCA or ADMIXTURE results. Samples clustered by geography rather than by batch (Fig. S8a–b), and there was no correlation between sequencing depth and ADMIXTURE proportions (Fig. S8c). We also did not observe significant effects of sequencing depth on PC1 or PC2, although PC3 and PC4 did show moderate correlations with depth (Fig. S8d). Overall these findings support the interpretation that the clustering patterns are predominantly biological rather than stemming from a technical artifact.

The chloroplast phylogeny also showed clear geographic structuring (Fig. S10). Of the major distinct populations identified in the ADMIXTURE and PCA nuclear SNP analyses, the Northern and Southeastern populations, in particular, are represented in well-supported cpDNA clades. The Southwestern population is found in two cpDNA clades, but these clades also have substantial membership from the Central population. The Central population, as assigned by nuclear SNPs, is found in every major cpDNA lineage, such as the clade containing samples from California and Oregon with only Central membership, which stems from a basal node of the tree. Samples assigned to the Northern population were the most cohesive in their phylogenetic clustering with the notable exception of Mystic 2, from the Black Hills of South Dakota, which fell within the main Southeastern chloroplast clade despite having 100% Northern nuclear ancestry (the three other samples from the Black Hills were in the Northern cpDNA clade). These patterns in the chloroplast, together, are thus largely consistent with the major nuclear lineages, but also suggest localized hybridization and chloroplast capture.

### Effective migration surfaces indicate elevated migration in the north

Results from FEEMS further emphasized a latitudinal trend in genetic structure. In the best-fitting model, effective migration was higher than average across the northern half of our sampling area (Fig. 2a). This area of higher effective migration roughly coincided with the distribution of the Northern population (Fig. 1). The southern half of our sampling range generally showed lower than average effective migration (Fig. 2a). FEEMS analysis of only the samples within this southern, historically unglaciated region revealed that the Southern Rockies had especially low average effective migration rates (Fig. 2b). Visualization of effective migration surfaces across varying levels of the penalization parameter *λ* further highlighted this pattern (Fig. S9): The Southern Rockies were notably the first feature to appear in the regularization path at high *λ* (Fig. S9a), indicating strong statistical support for this feature as a migration barrier (Marcus et al., 2021).

**Figure 2:**
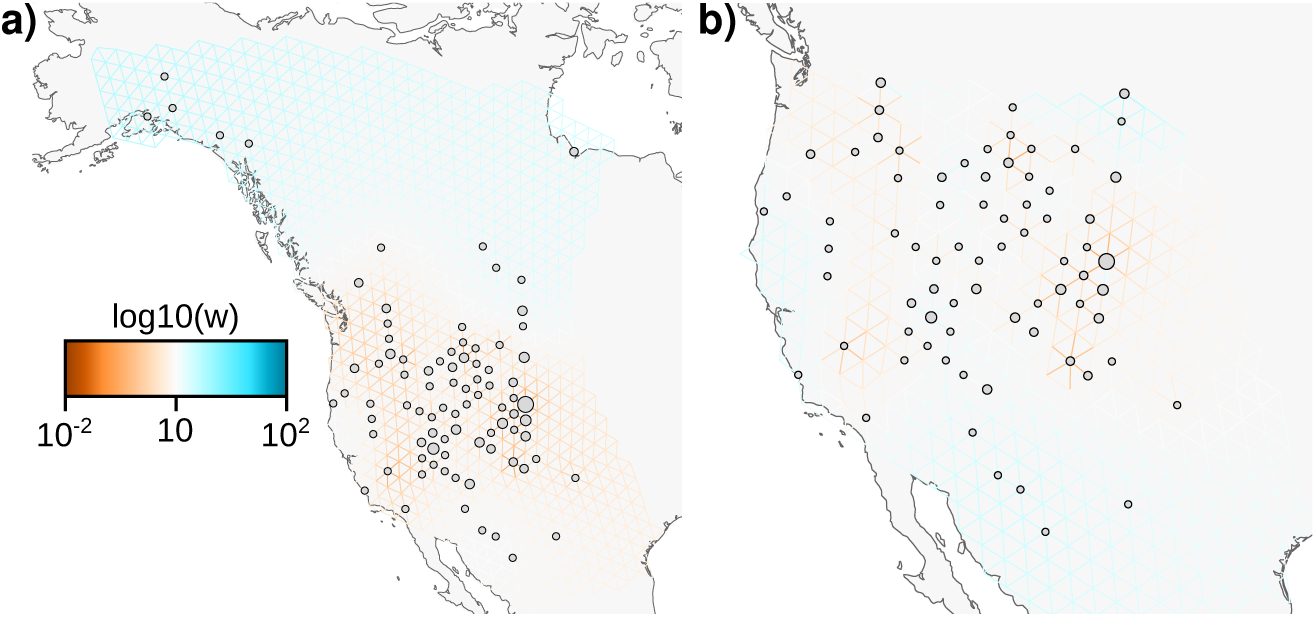
Lewis flax effective migration surfaces estimated with FEEMS for **a)** samples across its full range and **b)** its unglaciated range. Blue shading indicates higher than average effective migration; red shading indicates lower than average effective migration. Circles mark sample locations and circle size varies according to the number of individual plants assigned to the particular node on the grid.

### Genetic diversity is lower towards northern range margin

Nucleotide diversity (*π*) was highest in the Central population (4.46 × 10^−3^), intermediate in the Southwestern population (3.95 × 10^−3^), and lower in the Northern (3.55 × 10^−3^) and Southeastern (3.48 × 10^−3^) populations (Fig. 3). Continuous spatial windows of *π* clearly emphasized low diversity in northern regions (Fig. 3c. The low diversity of the Southeastern population as calculated with pixy was less clearly recapitulated in the wingen results, although a couple windows at the southern edge of our sampling did show reduced diversity (Fig. 3c). *F_ST_* was lowest for comparisons with the Central population (ranging from 0.096 to to 0.171) and highest between the Northern and Southeastern population (*F_ST_* = 0.242) (Fig. 3a). Absolute genetic divergence (*D_xy_*) ranged from 4.49 × 10^−3^ (N–C) to 4.81 × 10^−3^ (SE–C), indicating broadly similar absolute divergence among groups (Fig. 3a). In particular, the SE–N comparison had the highest *F_ST_* (0.242) but *D_xy_* was relatively moderate (4.65 × 10^−3^): low within-population diversity is the primary driver of higher *F_ST_* in this case. One possible explanation of the low nucleotide diversity within the SE populations is the more limited geographic area from which most of our collections of this group were made, compared to the other three populations: collections for the SE population were concentrated around Boulder County, Colorado. Given that we did not have *a priori* knowledge of geographic boundaries of genetic clusters, this was coincidental. Still, comparisons with the SE populations did have slightly higher *D_xy_* than those involving only the other three populations, suggesting the SE population is somewhat more differentiated. In all four populations, linkage disequilibrium decayed rapidly to background levels within 1 kb but was overall higher in the Northern population compared to the others (Fig. 3b). Inbreeding coefficients were also low, around zero, across all four populations (Fig. 3c). Within the Northern population, inbreeding coefficients showed a significant latitudinal cline, with plants sampled from more northern sites having higher *F* values (Fig. 3d). This trend was concordant with a northward reduction in observed heterozygosity among spatial windows encompassing the Northern cluster (Fig. 3e).

**Figure 3:**
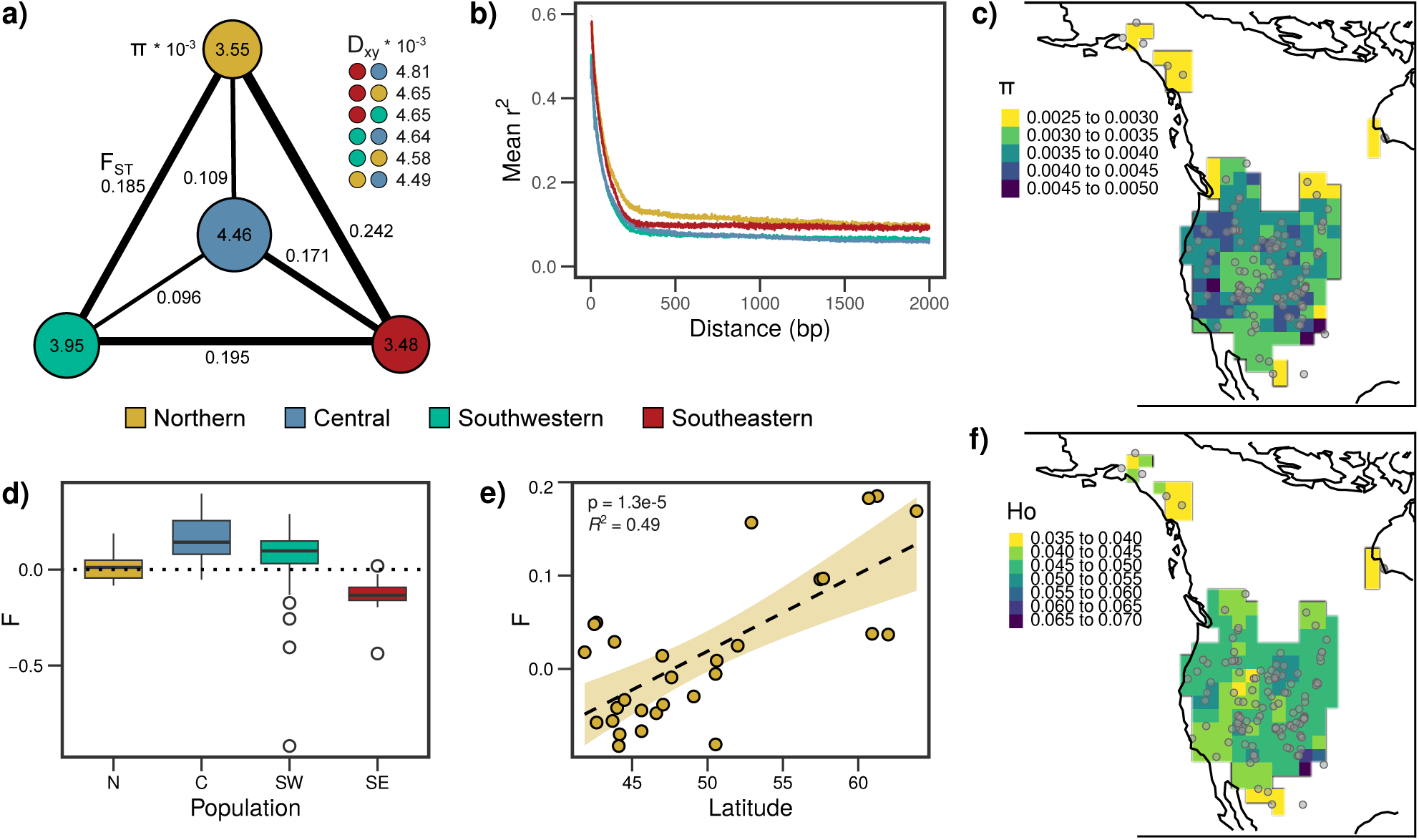
Population diversity, differentiation, linkage disequilibrium, and inbreeding statistics. **a)** Diagram representing population average pairwise nucleotide diversity (*π*) as circles and average pairwise *F_ST_* as edges. Circle size and edge thickness are proportional to the indicated values. Genome-wide average *D_xy_* for each population pair is shown in decreasing order. **b)** Linkage disequilibrium decay curves in the four populations. **c)** Nucleotide diversity (*π*) computed for moving windows across the landscape. **d)** Individual inbreeding coefficients (*F*) summarized at the population level. **e)** Inbreeding coefficients versus latitude within just the Northern population. **f)** Observed heterozygosity (Ho) averaged across individuals within moving spatial windows.

### Population size contractions and splits occurred during the Last Glacial Period

Modeling of effective population size (*N_e_*) through time with SMC++ revealed size contractions across all four populations that reached minima during the Last Glacial Period (115–11.7 kya) (Fig. 4). The Northern population showed the greatest contraction and lowest minimum *N_e_* (Fig. 4a). Timing of expansions in both the Central and Northern populations corresponded with the end of the Last Glacial Maximum around 20 kya, however the Northern population showed a much reduced expansion compared to the Central population (Fig. 4a). After the Last Glacial Maximum, the Southwestern population peaked at around 6 kya and subsequently contracted (Fig. 4a). All four populations had negative genome-wide Tajima’s *D* (N: -1.62, C: -2.37, SW: -1.96, SE: -1.26) (Fig. 4a), which corroborates the signals of recent expansion in all four populations.

**Figure 4:**
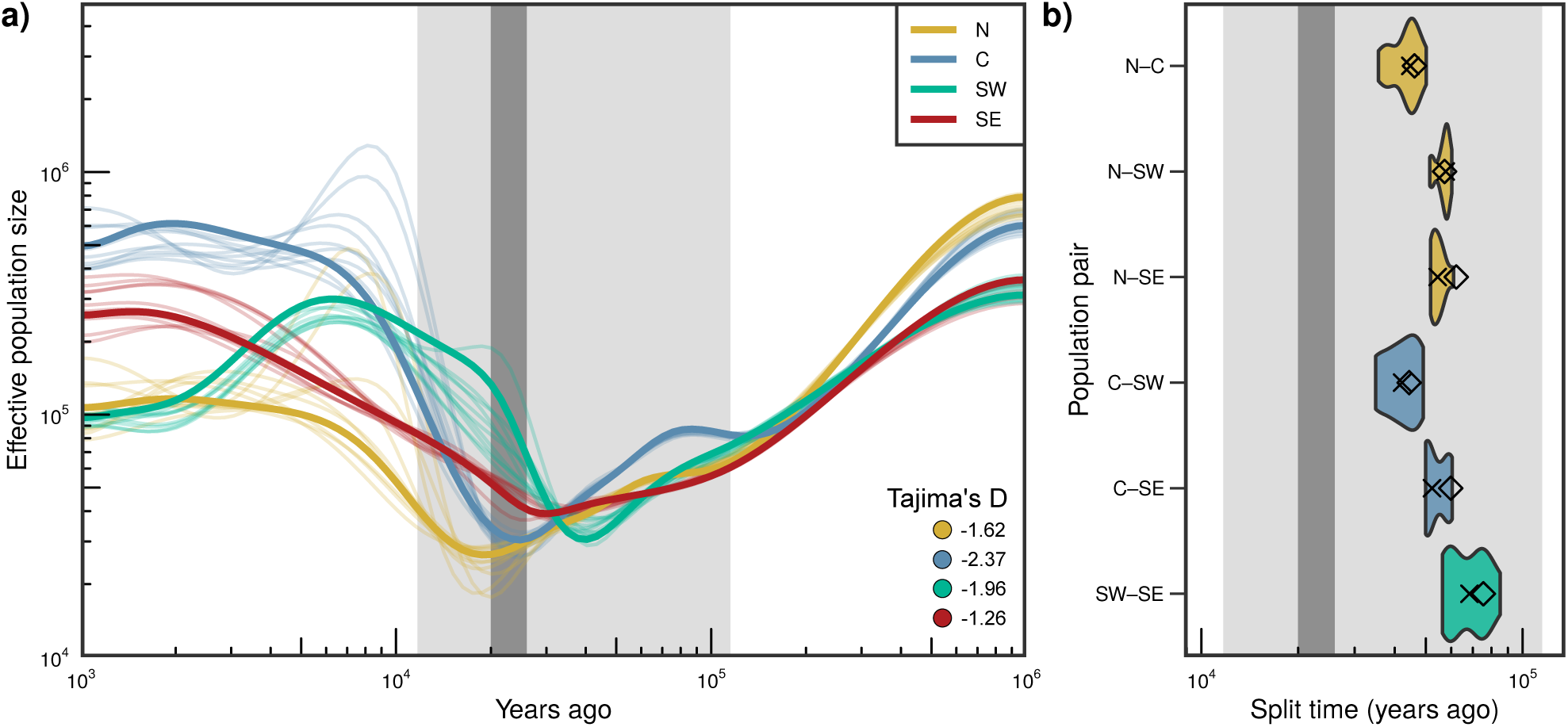
Demographic inference with SMC++ indicates population size contractions and splits during the Last Glacial Period. **a)** Population size histories 1,000–1,000,000 years ago. Thin lines represent 10 bootstrap replicates per population. Light gray and dark gray shading indicate the Last Glacial Period (115–11.7 kya) and Last Glacial Maximum (26–20 kya), respectively. Genome-wide average Tajima’s *D* for each population is also shown. **b)** Estimates of divergence times for each population pair. Diamonds indicate point estimates and violin plots represent bootstrap estimates with the median bootstrap value marked as a cross. Violin plots are colored according to the first population in the pair.

Divergence times for all population pairs were estimated to have occurred during the Last Glacial Period, before the Last Glacial Maximum (Fig. 4b). The oldest estimated split was between the Southwestern and Southeastern populations (median 68.4 kya). Split times were similar for Northern and Southeastern (median 54.4 kya) and Northern and Southwestern (median 57.6 kya) populations. Splits involving the Central population were estimated to have occurred most recently, ranging from 52.2 kya (C–SE) to 44.7 kya (N–C) and 42.1 kya (C–SW), which is consistent with these same pairs having the lowest *F_ST_* (Fig. 3a).

## Discussion

### Impacts of glaciation and topography on *Linum lewisii* population structure

Our results provide new understanding of major lineages within Lewis flax and point to the historical and contemporary factors that have shaped genetic variation in this widespread species. We found that the population genetic structure of Lewis flax has a clear geographic element best explained by the presence of four ancestral populations, named here as the Northern, Central, Southwestern, and Southeastern populations (Fig. 1). While we are unable to precisely pinpoint locations of glacial refugia, this structure is consistent with the “complexity” or *refugia within refugia* hypothesis (Gómez & Lunt, 2007; Shafer et al., 2010), in which multiple isolated refugia and historical substructure are present within, in this case, the classical southern refugium. This general trend has been frequently observed in North American plant and animal taxa, including rockcress (Song et al., 2006), western redcedar and western hemlock (Ruffley et al., 2022) gray fox (Kierepka et al., 2023), and black bear (Puckett et al., 2015).

Despite Lewis flax being a relatively cold-tolerant species (Gossweiler et al., 2024) with a hypothesized Beringian origin (McDill et al., 2009), evidence of a Beringian or other northern refugium was unapparent. While all populations showed evidence of recent expansion, the Northern population experienced the strongest bottleneck in genetic variation relative to historic levels, as well as the most recent expansion, at around the end of the Last Glacial Maximum (Fig. 4a). The Northern population showed a northward decline in genetic diversity that is consistent with postglacial northward range expansion from the south producing successive bottlenecks (Fig. 3e–f). Higher effective migration rates in the north versus lower migration in the south are also consistent with rapid postglacial northward expansion and southern refugia (Fig. 2). If Lewis flax did indeed colonize North America via Beringia, the current data suggest initial colonizing populations in the northwestern reaches of the continent did not persist through Pleistocene glacial cycles.

Likewise, significant genetic structuring along the latitudinal axis and an overall moderate association between genetic and geographic distance supports this scenario and indicates that isolation by distance contributes to Lewis flax population structure (Fig. S7). These results align with a trend of north–south genetic structuring in widely distributed western North American plant species, including yellow monkeyflower (Twyford et al., 2020) and quaking aspen (Callahan et al., 2013), as well as Great Plains species like common sunflower (Park & Burke, 2020), and silflower (Raduski et al., 2021).

Our results also emphasize a phylogeographic role for mountainous regions in western North America: the Central population was associated with both the Rocky Mountains and the Cascade–Sierra province, surrounding the Great Basin in a horseshoe fashion (Fig. 1). This is suggestive of mountainous regions acting as dispersal corridors for Lewis flax, as has been found for numerous species (Ren et al., 2025; Shafer et al., 2010; Tian et al., 2018). At the same time, we found that the Rocky Mountains mark west–east transitions from Central to Northern ancestry and from Central to Southeastern ancestry (Fig. 1), and effective migration surfaces point to the Continental Divide—particularly in the Southern Rocky Mountains—as a possible dispersal barrier (Figs. 2, S9). Regions where differentiated lineages meet violate the stepping-stone model that underlies FEEMS and thus may appear as migration barriers even under substantial contemporary gene flow (Al-Asadi et al., 2019; Petkova et al., 2016). Instead of being a persistent dispersal barrier, the Southern Rockies therefore could represent a zone of secondary contact between refugial populations separated by the mountain chain, with gene flow occurring during recolonization of high elevation habitats following glacial retreat (Jensen et al., 2024; Remington, 1968; Swenson & Howard, 2005). Little is known about Lewis flax seed dispersal except that like other flax species, its seeds form a sticky mucilaginous coating when wet. It’s unclear whether this provides a long-range dispersal mechanism via adherence to or ingestion by animals, or if it is primarily a means of anchoring to the ground (Kreitschitz et al., 2020; Yang et al., 2012).

Population divergence time estimates indicated an impact of the most recent glacial cycle on diversification in Lewis flax (Fig. 4). These estimates should be interpreted cautiously given sensitivity of the analysis to uncertainties of generation time, mutation rate, and post-split gene flow. One alternative scenario is that differentiation occurred along spatial and environmental gradients during postglacial range expansion from a single panmictic population. However, given that the clean-split model of SMC++ is likely to be biased towards more recent split times in the presence of post-split gene flow (Adrion et al., 2020; Terhorst et al., 2017), it is unlikely that we are overestimating the split times. Rather, our divergence time estimates are consistent with population divergence occurring as an ongoing process throughout the Pleistocene, with repeated cycles of isolation and contact eventually leading to differentiation. In this context the tight affinity of historical population sizes for C–N and SW–SE pairs *ca.* 1 Mya suggests shared history of these lineages and possibly a deeper split between more general northern and southern compartments predating the LGP (Fig. 4); this is consistent with C–N and SW–SE groups separating along the first principal component of SNP variation, as well as the topology of the neighbor-joining tree based on inferred allele frequencies of ancestral populations (Fig. 1), and the C–SE comparison having the greatest absolute genetic divergence (Fig. 3a). Additional demographic modeling that accounts for gene flow and the order of population splits, as well as hindcast niche modeling, could help clarify support for different phylogeographic scenarios. Furthermore, future work should explore patterns of adaptive genetic variation and the extent to which local adaptation has contributed to divergence among Lewis flax lineages, which we acknowledge as a ubiquitous evolutionary force.

### Taxonomy

Our findings regarding Lewis flax population structure do not neatly align with the current accepted infraspecific taxonomic treatment, which delineates three varieties. *Linum lewisii* var. *alpicola* is described as a high-elevation specialist inhabiting mountain ridges (2,000–3,700 m) in California, Idaho, Nevada, and Utah, with reduced stature, flower size, and style length (Flora of North America Editorial Committee, 2016). We sampled several plants from described *alpicola* habitat but did not observe any uniquely shared genetic ancestry among these samples (we note, however, that none of our samples were labeled *a priori* as var. *alpicola* and we did not assess phenotypes). Our sampling of Hudson Bay endemic *Linum lewisii* var. *lepagei* was limited to two herbarium specimens from a single area, both of which were grouped within the Northern population. In the chloroplast phylogeny, the *lepagei* samples were placed within a sub-clade of the main Northern clade alongside samples from Saskatchewan and North Dakota. The *lepagei* variety is thought to comprise a distinct lineage given its white flowers and separation from the core range (Flora of North America Editorial Committee, 2016; Mosquin, 1971), however, this was not apparent in our current sampling. Our analyses of genetic diversity and population demographic history support the idea that Lewis flax recolonized northern latitudes recently, following the Last Glacial Maximum (Figs. 3, 4). Therefore, it’s plausible there has not been sufficient time for the *lepagei* variety to clearly differentiate from the rest of the Northern population. Improved sampling of var. *lepagei* as well as var. *alpicola* would be critical to make more conclusive determinations regarding their origins and genetic variation.

The third currently accepted infraspecific, *L. lewisii* var. *lewisii* encompasses the four main populations described here. In our experience, these populations do not show obvious, discrete phenotypic differences, but there is some evidence from previous crossing experiments that reproductive barriers exist between them: hybrids between an Alaskan population (i.e. Northern) and Texan population (putatively Southeastern) showed diminished pollen viability compared to hybrids between the same Texan population and a USDA collection of uncertain origin but “probably from the western United States” (Mosquin, 1971). Future work should explore such crossing barriers more extensively and could lend support to updated taxonomic treatments more in line with population genetic structure.

### Mating system evolution in North American blue flax

Our results indicate that Lewis flax is predominantly outcrossing despite its loss of distyly and shift to homostyly and self-compatibility. For instance, inbreeding coefficients overall were close to zero and linkage disequilibrium decayed rapidly (Fig. 3). Effective population sizes in recent history were also moderate to high (Fig. 4). These findings give support and new context to previous pollination experiments showing Lewis flax of central Colorado origin does not set seed in the absence of insect pollinators (Kearns & Inouye, 1994). Predominant outcrossing is also harmonious with its perennial habit, as strong selfers tend to be annual. Thus, Lewis flax provides a notable counter to the expectation of distyly loss leading to predominant selfing (Gutiérrez-Valencia et al., 2024), although it is likely not unique in this regard: recent study of its close relative *Linum leonii* also revealed evolution of mixed mating following distyly loss (Postel et al., 2026).

The latitudinal cline in inbreeding coefficients within the Northern population (Fig. 3e) was an intriguing result in light of past observations of reduced herkogamy in northern plants (Mosquin, 1971). While our individual-based sampling scheme means we cannot fully eliminate potential bias resulting from local allele frequencies differing from the cluster average when calculating inbreeding coefficients (i.e. Wahlund effect), we also observed reduced individual-based heterozygosity rates (*H_o_*) in northern areas (Fig. 3f), indicating that the inbreeding cline is not artifactual. These patterns in the Northern cluster—in addition to its elevated linkage disequilibrium (Fig. 3b) and lower *N_e_*, (Fig. 4a)—could simply be the result of smaller, patchier, and more bottlenecked populations that established in formerly glaciated terrain. However, they are also consistent with northern plants having higher selfing rates, supported by observations of reduced herkogamy in northern plants (Mosquin, 1971) and the general link between herkogamy and outcrossing–selfing (Opedal, 2018), including the fact that strong selfing is linked to diminished herkogamy in Lewis flax’s sister taxa, *Linum pratense* (Uno, 1984). More intensive population-level sampling coupled with measurements of herkogamy will be needed to test this hypothesis and clarify the link between postglaciation range expansion and increased selfing rates in Lewis flax, although this scenario has been documented in other widespread North American forbs, such as American bellflower *Campanula americana* (Koski et al., 2019). We note, however, that even with possible increased selfing rates, Northern plants retain substantial heterozygosity and genetic variation. In sum, Lewis flax may have a “best of both worlds” reproductive strategy—the ability to self provides reproductive assurance, while predominant outcrossing limits inbreeding depression (Goodwillie et al., 2005). We speculate that this contributed to its colonization of North America as well as its broad prevalence on the continent.

### Implications for restoration and agriculture

Lewis flax is a popular species included in seed mixes for restoration of native plant communities, especially on disturbed lands in the Intermountain West (Kitchen, 1995; Leger & Baughman, 2015; Winslow et al., 2009). Our study provides a foundation to guide land managers in making decisions about seed sourcing and transfer. Currently, these decisions are complicated by long-standing concern of hybridization between Lewis flax and its introduced congener, *Linum perenne*, which is native to Eurasia. In the 1980s, the United States Department of Agriculture (USDA) released what was claimed to be a native blue flax cultivar, “Appar”, but which turned out to be a collection of *Linum perenne* (Pendleton et al., 2008b, 2008a). Its subsequent and ongoing use in restoration and horticulture has led to it being widespread and naturalized across much of western North America [Pendleton et al. (2008b); Pamela Cornelisse, Christina Littleton, Becky Pierce pers. comm.]. Our study provides necessary genomic resources to test whether populations of interest are “pure” *L. lewsii* genotypes suitable for future development into new seed sources to enhance landscape management efforts across western North America. Some preliminary evidence suggests admixture between native and introduced blue flax is largely absent, even in places where the two species occur in close proximity, for instance in Boulder County, Colorado (Innes, 2024).

The USDA rectified the mistaken identity of Appar in 2003 upon selection and release of the “Maple Grove” Lewis flax pre-variety germplasm in 2003, which was originally collected from Millard County, Utah. Maple Grove seed is commercially grown and widely available from seed companies and garden stores, and commercial production of Lewis flax seed generally has focused on populations from the Great Basin due to ecosystem restoration efforts in this region creating high demand for native seed (Innes et al., 2022; Pendleton et al., 2008a). Human-facilitated migration could therefore explain the disjunct presence of Southwestern-type samples far outside the main geographic region of that population (Fig. 1). This is most assuredly the case for the accession from an isolated population in the Appalachian mountains of West Virginia (Ames 34946, Table S1). It is plausible that humans also moved another accession of Southwestern ancestry, originally collected from a roadside area in northern Wyoming (Ames 34266), as well as three samples of Southwestern ancestry collected in central Colorado (Guanella Pass High, Dow Mountain, and Homestake Reservoir). The aforementioned samples were assigned 100% Southwestern ancestry despite being geographically distant from the southwestern region and were all collected after the 2003 commercial release of Maple Grove. Homestake Reservoir had the same chloroplast haplotype as Maple Grove, while the other disjunct Southwestern-type samples clustered with those from Nevada (Fig. S10). This suggests that Southwestern genotypes, including Maple Grove, have been moved far beyond their original provenance.

Increasing human-mediated dispersal and prevalence of Southwestern genotypes could have consequences for population structure and evolution of Lewis flax into the future. Maple Grove or other Southwestern genotypes might not be suitable in particular regions if genetic differentiation has involved local adaptation, or if maintaining genetically distinct local populations is a priority. These considerations underlie traditional seed sourcing strategies aiming for regionally-sourced, locally-adapted seed (McKay et al., 2005). On the other hand, newer restoration strategies, such as climate-adjusted provenancing, seek to account for a rapidly changing climate by combining local seeds with those from historically hotter and drier sites to better match climate predictions at target sites (Prober et al., 2015). With this approach, the Southwestern gene pool may be useful for seed transfer into other regions. Either way, we encourage land managers to take into account the spatial population structure described here when sourcing Lewis flax for restoration projects.

There is also an ongoing effort to domesticate Lewis flax as a perennial oilseed crop in part due to its seeds being rich in healthy *α*-linolenic acid (Innes et al., 2022; Pull et al., 2023). The results we present here can help guide breeding and future collection efforts (Raduski et al., 2021). The major genetic lineages we identify here may not be easily intercrossed (Mosquin, 1971) and additional data from crossing experiments are needed to better understand these incompatibilities. Such crossing barriers may make trait transfer from one genetic lineage to another via breeding difficult or impossible. Additionally, future seed collection efforts could be targeted to areas where limited seed is available in current repositories, particularly the southern Great Plains, California, Mexico, and Canada, where our sampling mainly came from herbaria if at all. This work also will inform development of diversity panels and novel study populations (such as nested association mapping panels) for future trait genetic mapping.

## Conclusion

Diversification of *L. lewisii* across its range was previously not fully examined genetically, and here we have described its population structure comprising four main genetic lineages shaped by late Pleistocene glaciation. This structure is largely cryptic because these lineages do not correspond with current infraspecific taxonomy or obvious phenotypic differences. These findings align well with an observation made decades ago that in Lewis flax “the buildup of the kind of reproductive barriers that lead to the sterility of interpopulational hybrids is a process that has proceeded more or less independently of gross morphological differentiation” (Mosquin, 1971). However, a thorough examination of phenotypes might find traits that do align well with the major genetic lineages within the species.

Another enigmatic aspect of *L. lewisii* is its approach herkogamy, which is anomalous among homostylous, self-compatible blue flaxes. We have shown that, despite its shift to self-compatibility, Lewis flax exhibits genomic signatures suggestive of predominant outcrossing. Thus, loss of distyly does not necessitate a shift to predominant selfing. Future work investigating diversification of North American blue flax should also consider *Linum pratense*, which exhibits a more conspicuous morphological selfing syndrome but is poorly characterized from a genetic and taxonomic standpoint.

On the other hand, *L. lewisii* is exemplary of a number of recurrent North American phylogeographic trends. Its population structure and demographic history suggest the existence of multiple glacial refugia existing south of the continental ice sheet, and it displays a genetic diversity cline indicative of a classic postglacial northward range expansion; a more general trend of north–south genetic structuring, frequent in North American taxa, is also present in Lewis flax. Lastly, our genomic analyses emphasize the Rocky Mountains as a multi-faceted evolutionary force, likely as a barrier separating distinct refugia during glacial periods and as a contact zone and dispersal corridor during interglacial periods and following the Last Glacial Maximum. Overall, this study illustrates the genetic legacy of the last Ice Age in North America and contributes to our understanding of the relationship between continental-scale geography and mixed mating systems in driving diversification.

## Data availability

Raw sequence data generated and used in this study are publicly available on NCBI Sequence Read Archive under BioProject PRJNA1306336.

## Author contributions

PI: Conceptualization, Methodology, Formal analysis, Investigation, Writing - Original Draft, Writing - review & editing, Visualization, Funding acquisition, Project administration. ZM: Methodology, Investigation. BSH: Conceptualization, Investigation, Writing - review & editing, Resources. NCK: Conceptualization, Methodology, Writing - review & editing, Supervision, Resources, Funding acquisition.

## Acknowledgments

The authors kindly thank the following herbarium curators and collection managers for obliging our sampling requests: Burrell E. Nelson at Rocky Mountain Herbarium (RM), Diana Sawatzky and Bruce Ford at University of Manitoba Herbarium (WIN), Andrew Salywon at Desert Botanical Garden (DES), Massimo Boscolo at Arizona-Sonora Desert Museum (ASDM), and Dina Clark and Erin Manzitto-Tripp at University of Colorado Herbarium (COLO). The authors thank David Tork and Neil Anderson for providing plant tissue from multiple flax seed bank accessions, Brian Smart for greenhouse help, André Gossweiler for help with field collections, and Dan Klee for helpful discussion of phylogenetic methods. We additionally thank members of the Kern–Ralph co-lab for their feedback and suggestions.

