## Supplemental Materials for "Glaciation history and geography shape genetic diversity of Lewis flax (*Linum lewisii* Pursh.) across North America"

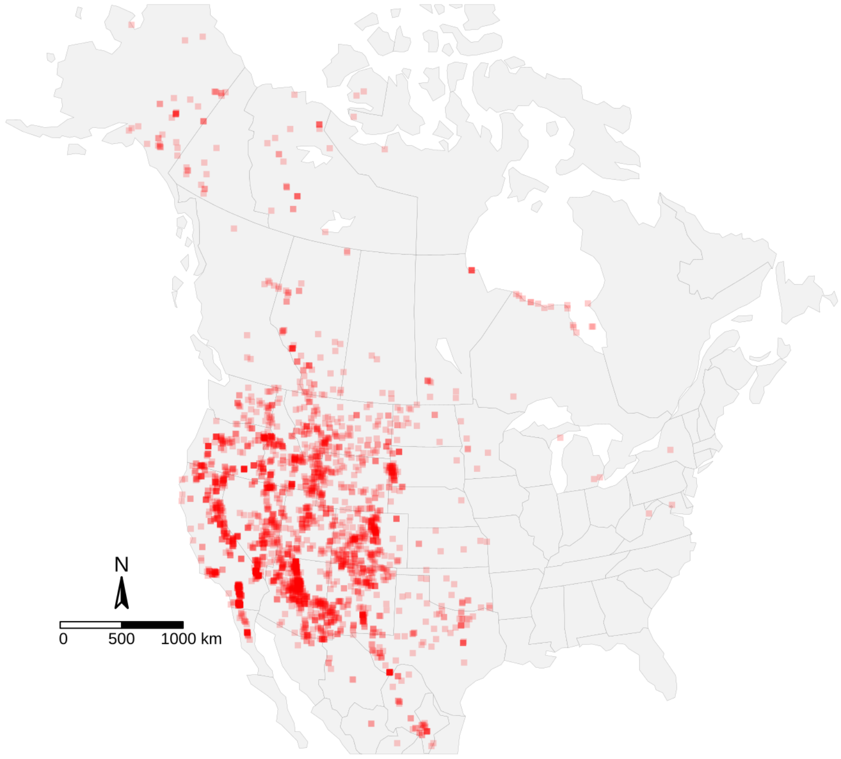


Figure S1: Lewis flax occurrences based on preserved specimen records. We downloaded occurrence data from iDigBio with a search for taxa names *Linum lewisii*, *Linum lewisii* var. *lewisii*, *Linum lewisii* var. *lepagei*, *Linum lewisii* var. *alpicola*, *Linum perenne var. lewisii*, and *Adenolinum lewisii*, which resulted in 2,813 occurrences.

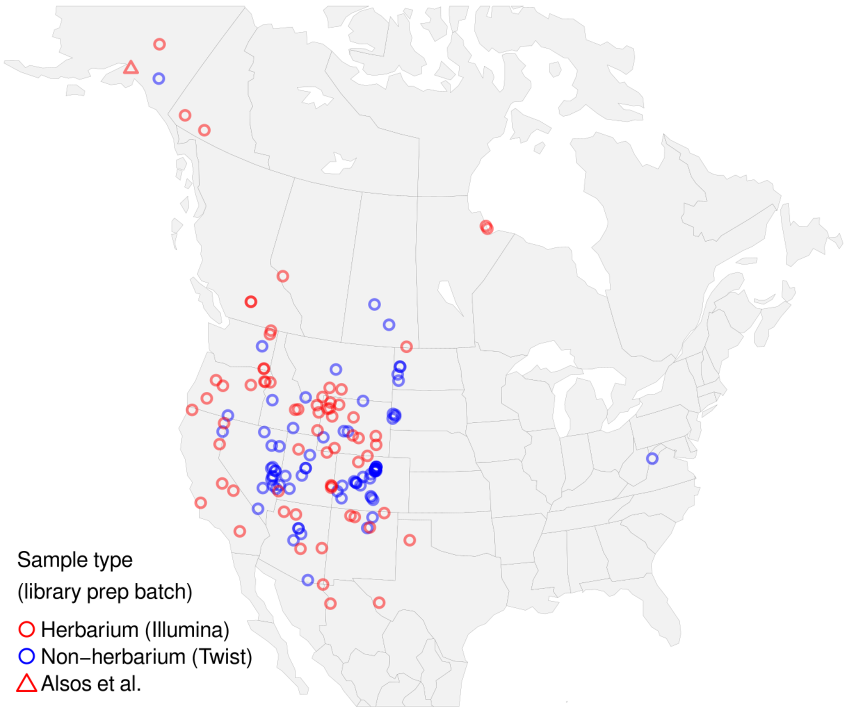


Figure S2: Locations of samples included in this study.

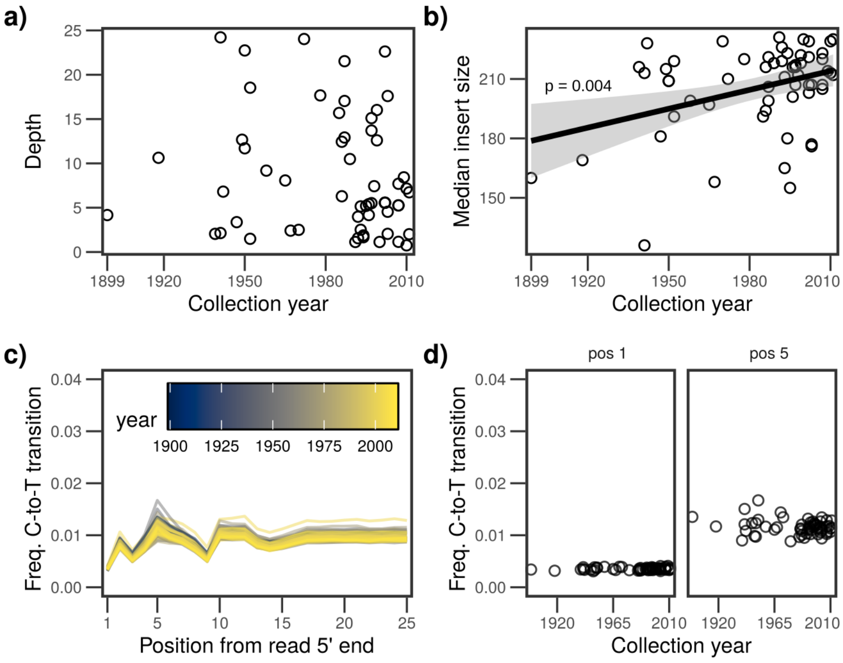


Figure S3: Herbarium DNA quality assessments. **a)** Per-sample average sequencing depth versus specimen collection year. **b)** Median insert size of sequence libraries versus specimen collection year **c)** Frequency of C–T transitions (i.e. cytosine deamination) at 5' end of reads as measured with mapDamage2. **d)** Frequency of C–T transitions at positions 1 and 5 at 5' read ends, versus specimen collection year.

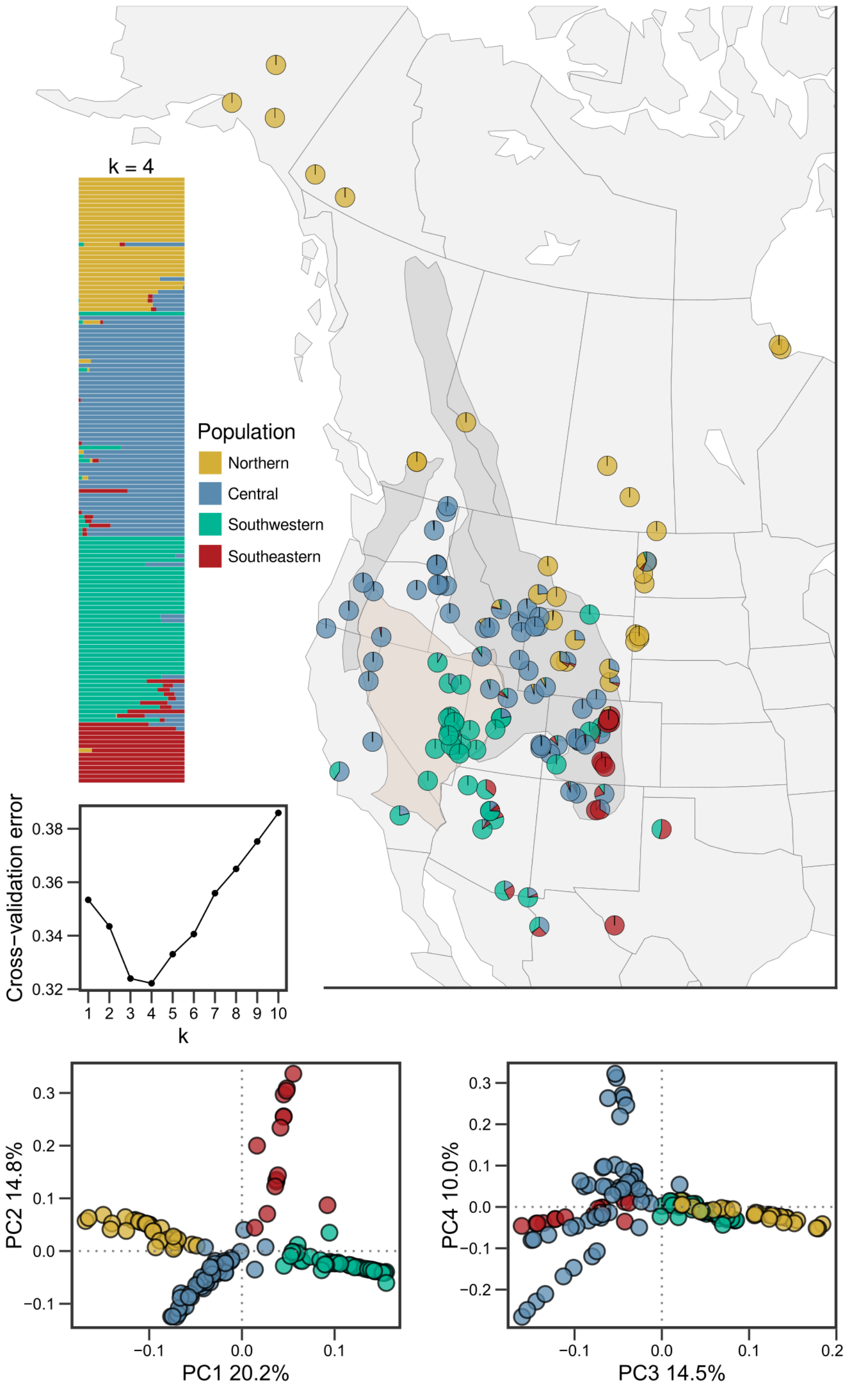


Figure S4: ADMIXTURE and principal component analysis results after excluding 20 of 28 Boulder County samples (reduced cohort). The darker gray shading represents approximate boundaries of the Rocky Mountains and Cascade Range, and the beige shading is the Great Basin. Sample points in PCA plots are colored based on the relative majority ancestry coefficient from the reduced cohort ADMIXTURE analysis at $k = 4$.

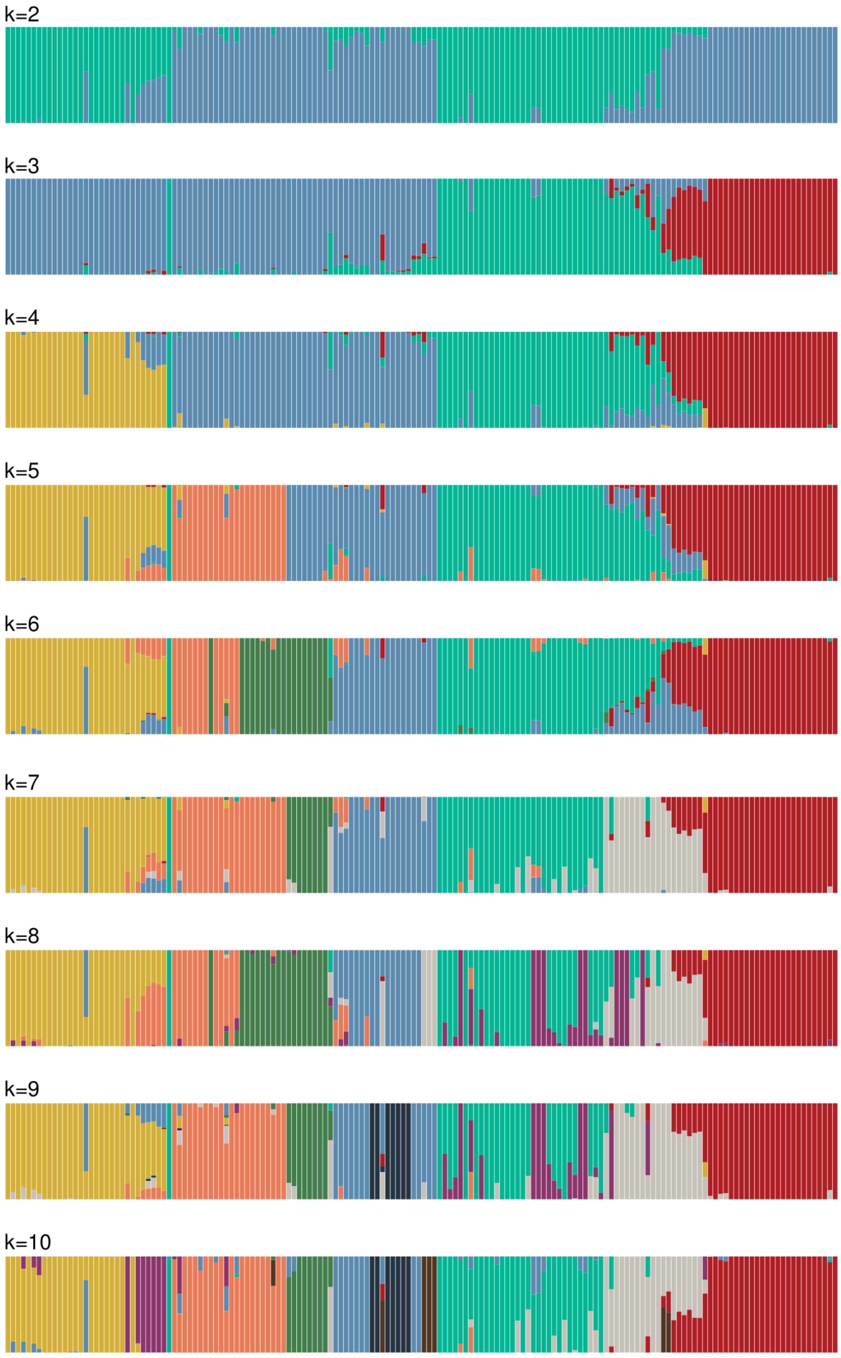

Figure S5: ADMIXTURE results for the full cohort with $k = 2$ through $k = 10$.

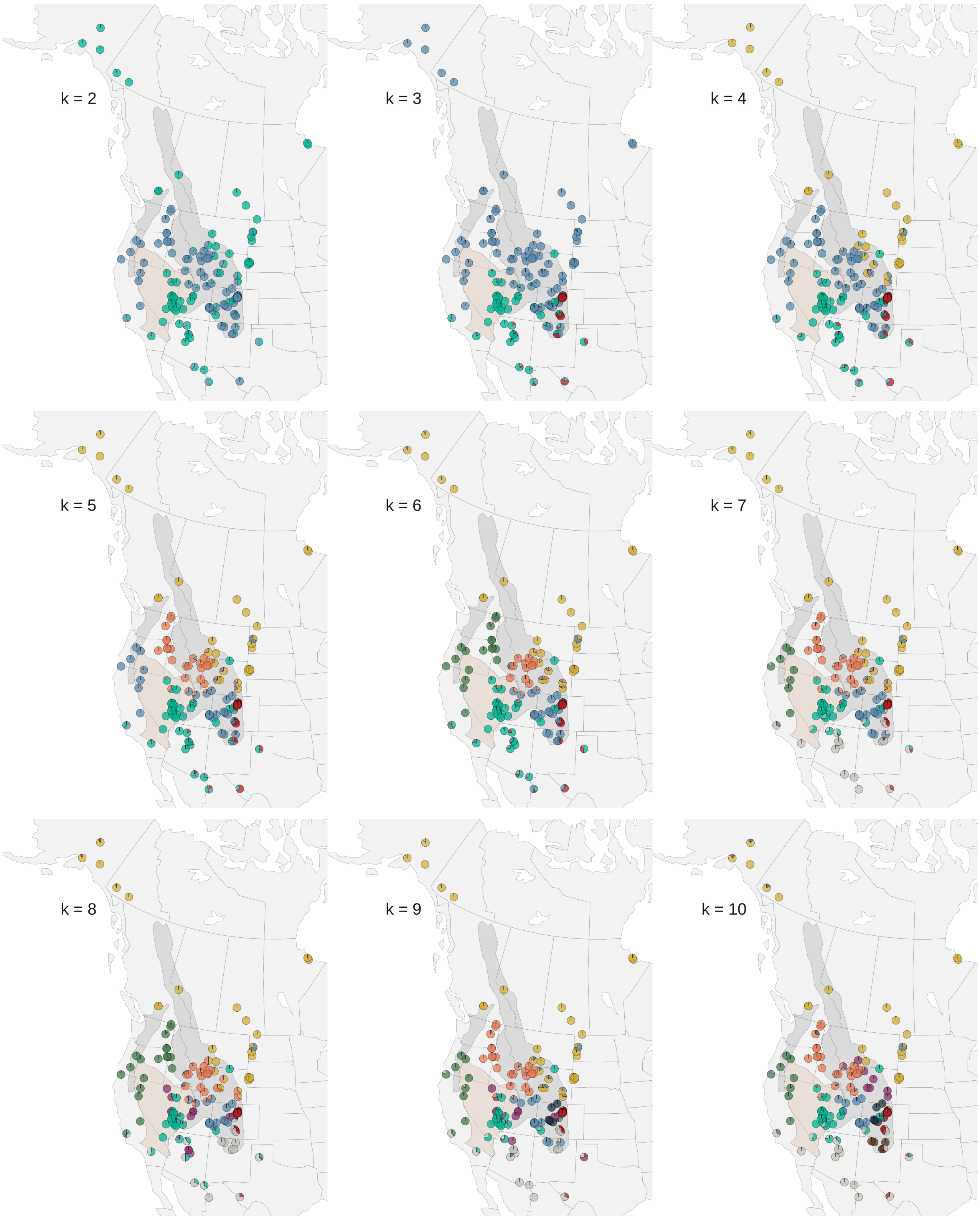


Figure S6: ADMIXTURE results visualized geographically for the full cohort with $k = 2$ through $k = 10$.

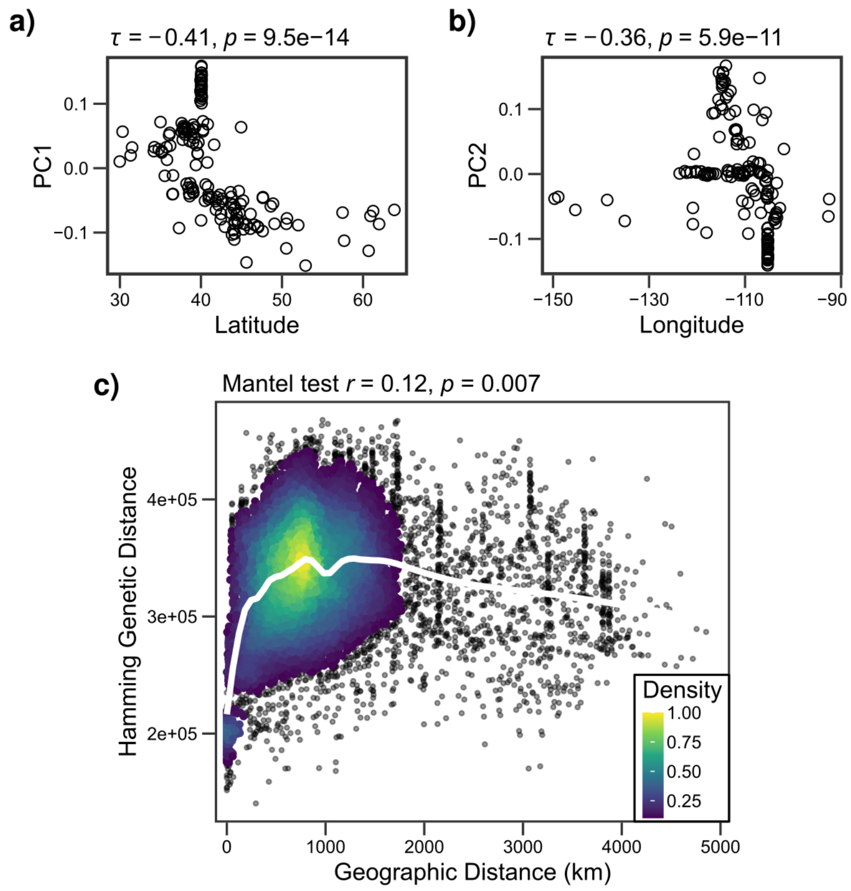


Figure S7: Geographic clines in principal components 1 **(a)** and 2 **(b)** with corresponding rank correlation coefficients (Kendall's tau) and p-values. **c)** Sample-pairwise geographic and genetic distances.

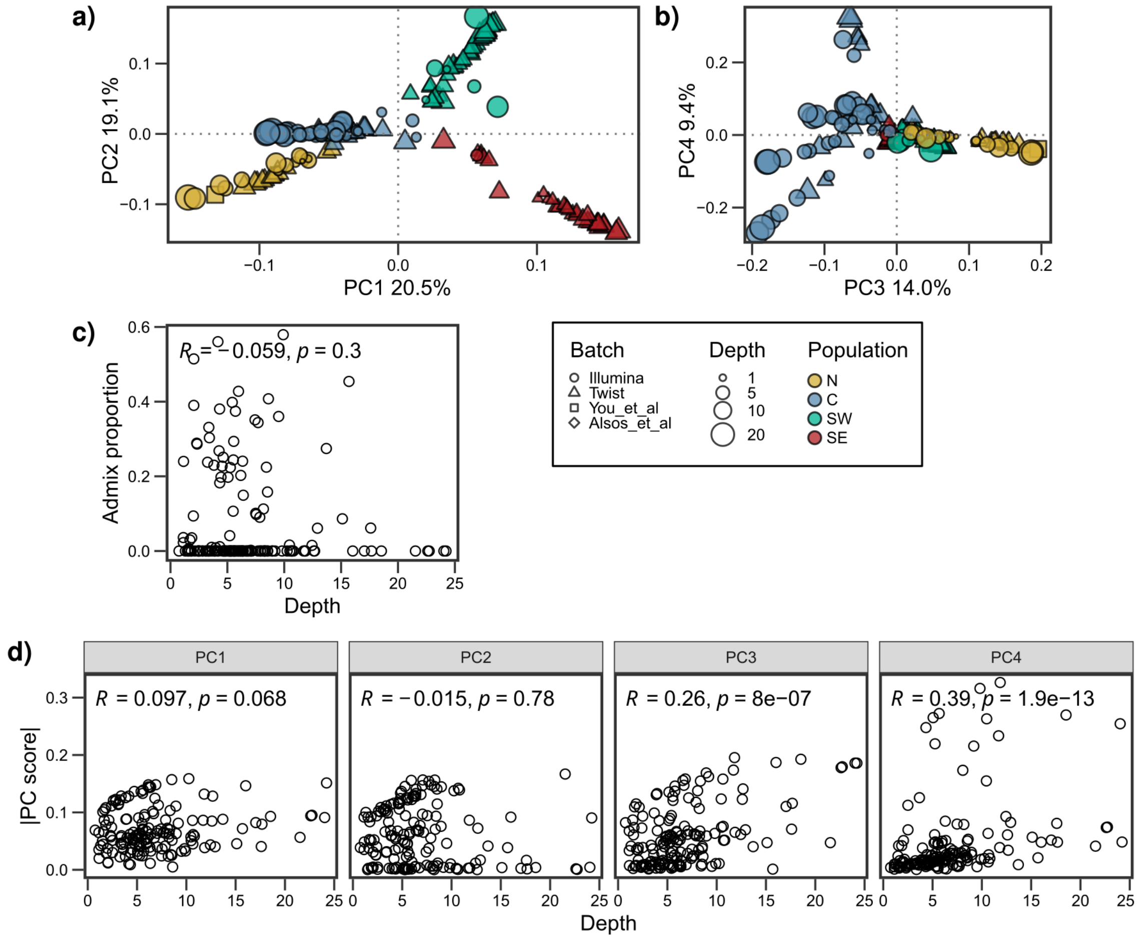


Figure S8: Effects of sequencing depth and batch on population structure inferences. **a–b)** PCA results as shown in the main text but with sequencing depth and batch represented by size and shape, respectively. Illumina and Twist batches correspond to the herbarium and non-herbarium samples, respectively, and refer to the kits used for their DNA library preparations. "You et al" and "Alsos et al" specify individual samples from previous studies. **c)** Effect of sequencing depth on admixture proportion as measured by the rank correlation coefficient (Kendall's tau). **d)** Effect of sequencing depth on principal components 1–4 as measured by Kendall's tau.

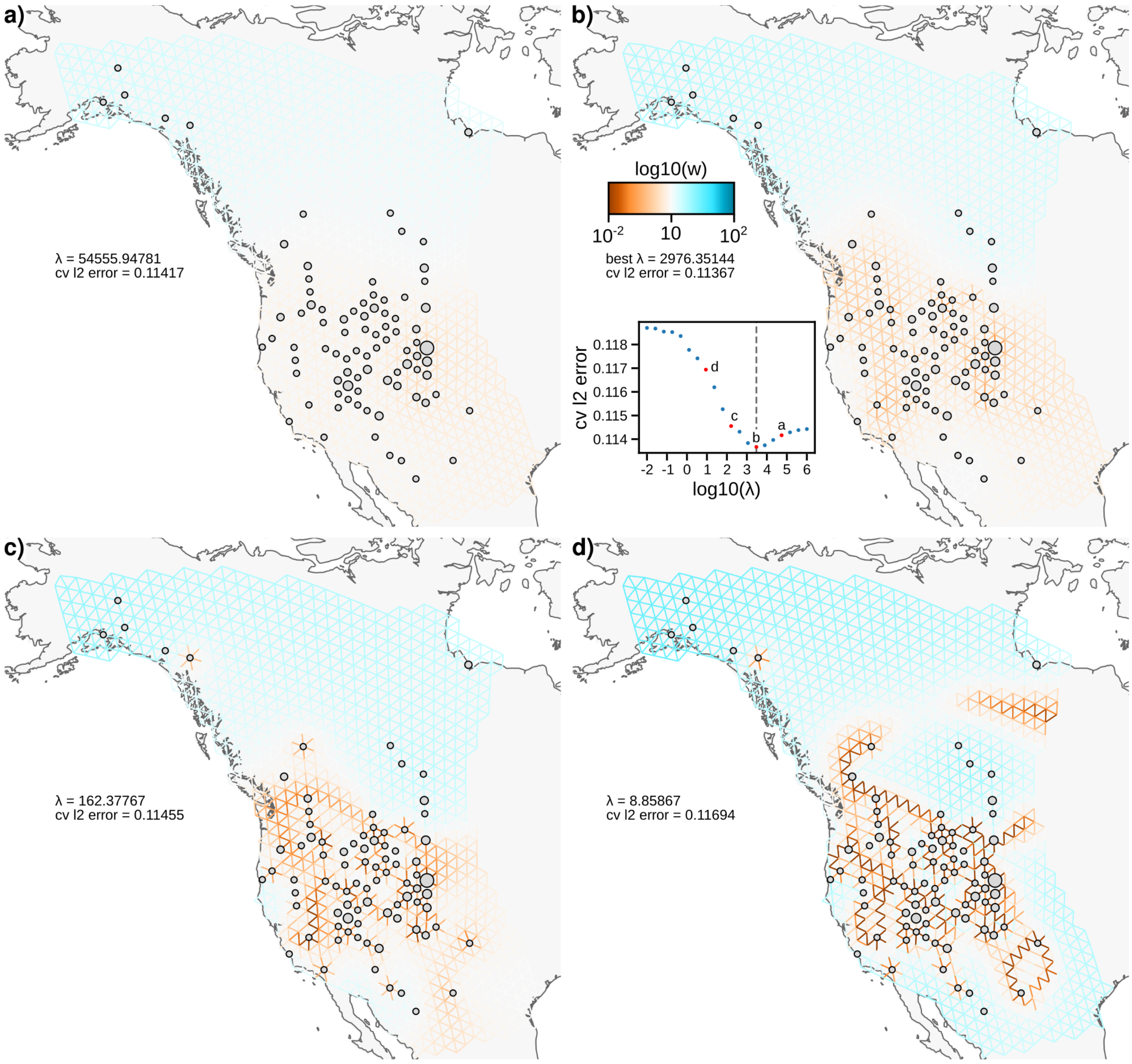


Figure S9: FEEMS results across varying values of $\lambda$. Circles represent sample locations, and circle size varies according to the number of individual plants assigned to the particular node on the grid. Panel **a)** shows that even when $\lambda$ is high, i.e. when there is a strong penalty for heterogeneous migration, the Southern Rocky Mountains have lower than average effective migration. Panel **b)** represents the iteration with the lowest cross-validation error and is the same as presented in the main text. Lower levels of $\lambda$, e.g. panels **c–d**, allow for more flexible models that may represent local structure but at the risk of over-fitting.

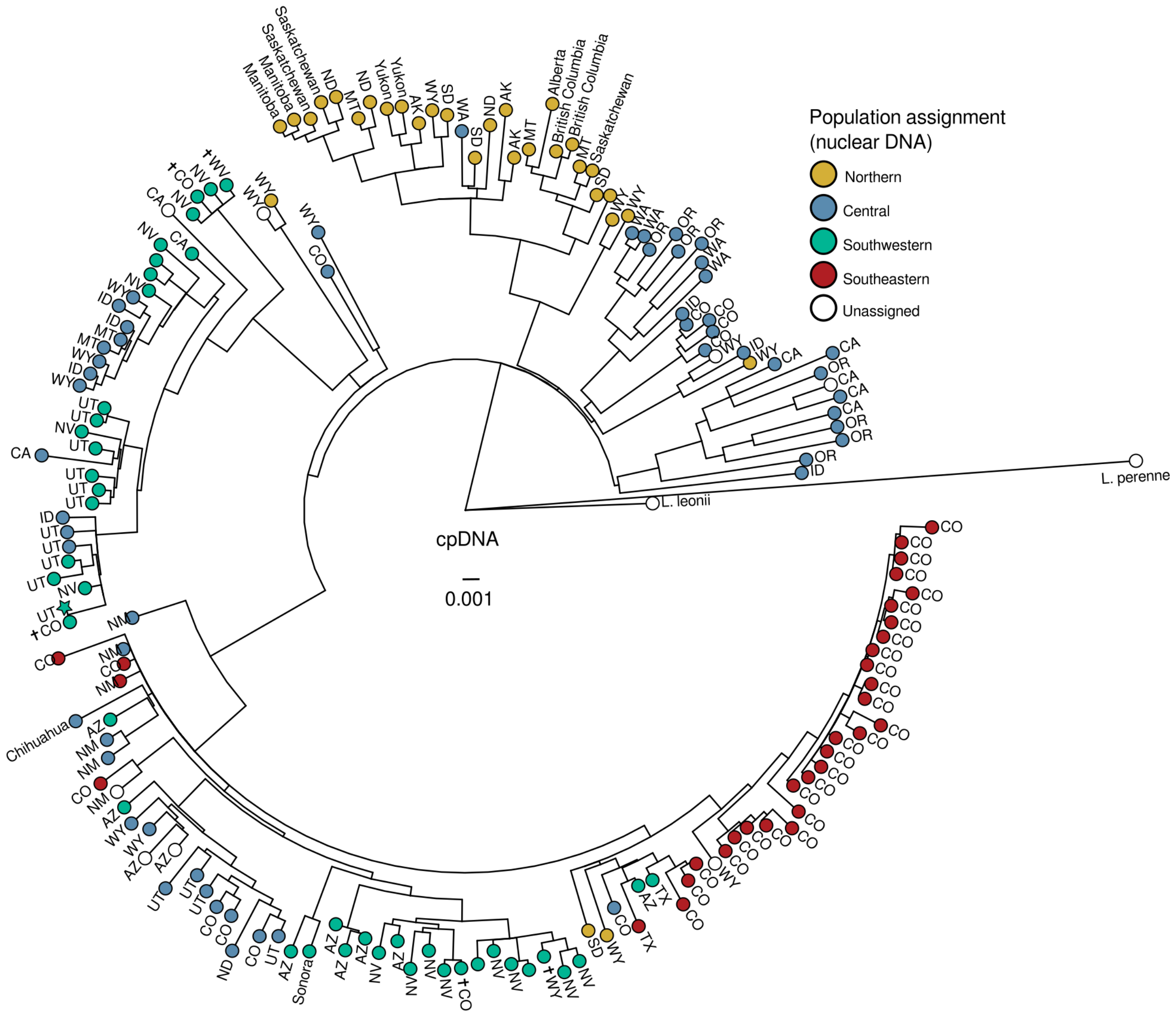


Figure S10: Maximum likelihood phylogeny inferred from whole chloroplast SNPs. Colors represent population assignment based on nuclear genome ancestry proportions as shown in the main text. Tips are labeled with sample state or province of origin. Maple Grove, the reference genotype, is shown as a star. Latin cross symbols indicate the disjunct Southwestern-type samples that were collected from far outside the core southwestern region. Empty (white) circles indicate the outgroup samples and Lewis flax samples which were excluded from analyses of nuclear genome variation due to low sequencing depth and thus lack a population assignment. Nodes with bootstrap support less than 75 are collapsed.
